# Selective genetic targeting of the mouse efferent vestibular nucleus identifies monosynaptic inputs and indicates function as multimodal integrator

**DOI:** 10.1101/2025.09.15.676291

**Authors:** Miranda A. Mathews, Victoria W. K. Tung, Emily Reader-Harris, Andrew J. Murray

## Abstract

The vestibular system is a critical sensory modality required for coordinated movement, balance and our ability to interact with the surrounding environment. Vestibular sensory neurons provide the nervous system with information about head rotation and acceleration. However, the nervous system can also modify the activity of sensory neurons and hair cells via the actions of the efferent vestibular system (EVS). The function of the EVS has remained unknown partly because of an inability to target efferent vestibular neurons in a selective manner to understand their synaptic inputs and function during behaviour. Here, we present a novel method for the selective targeting and expression of flp-recombinase in EVS neurons. We take advantage of the dual expression of choline acetyl transferase (ChAT) and calcitonin gene related peptide (CGRP) in these neurons to develop an adeno-associate virus (AAV) that expresses a gene only in neurons with this intersectional expression. We use this system to map the monosynaptic inputs to EVS neurons and show inputs from distinct populations of brainstem and midbrain regions indicating a functional role as a multimodal processing center and integrator for the vestibular periphery. To demonstrate the applicability of our technology in behavioural assays, we performed a preliminary behaviour analysis in mice with disrupted EVS function. While more bespoke assays are required to ascertain EVS function/s, our viral method presents a novel tool for investigators examining the role of the vestibular system and its central circuits.

## Introduction

The vestibular system is indispensable for the coordination of movement, balance and our general ability to interact with the environment. This ‘sixth sense’ provides information regarding head rotation and acceleration, which is combined in the brain with other sensory modalities to give a sense of the position and movement of the body in space.

Anatomically, vestibular organs for all land vertebrates are in the inner ear and vestibular sensory neurons relay information to the brainstem. Aquatic vertebrates possess an additional system, the lateral line, that detects changes to the surrounding aquatic environment (such as pressure, vibration, movement) and is critical for orientation underwater. Like several sensory systems communication is bidirectional, where the central nervous system can exert influence over the peripheral sensory organs. A central modulation of vestibular end organs has been observed across all vertebrates – and is termed the efferent vestibular system (EVS). The EVS originates in the efferent vestibular nucleus (EVN) of the brainstem and EVN neurons terminate directly on vestibular hair cells and afferent sensory neurons. In amphibians, efferent fibres innervate both the inner ear and lateral line (Hellmann and Fritzsch, 1996). However, while we know the EVS has specific cellular actions on vestibular and lateral line sensory receptors (for reviews, see Mathews et al., 2017; Holt et al., 2010), a specific behavioural and functional role is yet to be understood.

Over the last seven decades, researchers have adapted a variety of *in vivo* and *in vitro* techniques to understand the function of this small and evolutionarily conserved nucleus. These experiments fall into three broad categories: 1) EVS stimulation (e.g. chemically, electrically or thermally (most recently in Raghu et al., 2019) combined with recording peripheral vestibular activity (summarised in Mathews et al., 2017); 2) putative physiological EVN recordings under different behavioural paradigms (some examples include, Russell, 1971; Highstein and Baker, 1985; Plotnik et al., 2002; 2005; Chagnaud et al., 2015); and 3) mouse transgenic lines combined with behavioural recordings (Luebke et al., 2014; Hübner et al., 2015; 2017; Chang et al., 2021). Collectively, these works give insights into the potential function of the EVS (for reviews, see Holt et al., 2010; Mathews et al., 2017; Mathews et al., 2020; Poppi et al., 2020; Cullen and Wei, 2021), especially when paired with direct physiological recordings of EVN neurons (Leijon and Magnusson, 2014; Mathews et al., 2015). However, a complete understanding of EVS function requires an ability to target and manipulate EVN neurons in awake behaving animals – which so far has not been possible.

Recombinase mouse lines, where cre or flp is expressed in genetically defined neurons, have revolutionised our understanding of behavioural roles of neuronal subtypes. In the EVN, which is a cholinergic nucleus, Lorincz et al., 2021 utilized a ChAT-Cre mouse line to perform anatomical tracing of EVS neurons. However, cholinergic neurons are widespread in the brain and perform multiple functions so behavioural assessment of EVS function requires a finer scale of neuronal subtype resolution. Similarly, short enhancer or promoter sequences can be used within viral vectors such as adeno-associated virus (AAV) to drive cell-type-specific gene expression. In addition to ChAT, calcitonin gene related peptide (CGRP) is known to be expressed in EVN neurons (Mathews et al., 2015). Several groups have described CGRP promoter fragments for use in sensory or central neurons (Watson et al., 1995; Messina et al., 2000; Durham et al., 2004; Shimizu et al., 2017), providing an opportunity to utilise these same sequences for EVN targeting.

Here, we combine the ChAT-Cre transgenic mouse line with both cre-dependent AAVs and a cell-type-specific short promoter to intersectionally target brainstem neurons expressing both ChAT and calcitonin gene related peptide (CGRP). This intersection permits selective targeting of EVN neurons, which co-express both ChAT and CGRP (Ohno et al., 1991; Mathews et al., 2015). We first validated this approach by retrograde labelling of EVN neurons by applying fluoro-gold to the end organs. Following this validation, we were able to use this novel approach to identify the monosynaptic inputs to EVN neurons using modified rabies virus (RABV) and separately block neurotransmitter release from the EVN through the directed expression of tetanus toxin light chain (TeLC). This approach therefore allows us and others to elucidate the role of the EVS in gross motor and vestibular coordination.

## Materials and Methods

### Generation of AAV constructs

AAV constructs using the CGRP short promoter were based on a promoter sequence from Durham et al. (2004) and Thomas et al. (2001). We de novo synthesised (GeneArt, Life Technologies) a 2Kb region immediately upstream from the CGRP-β coding region. The sequence of the promoter region is as follows:

5’

ACGCGTGGTACCCCTGCCAGCTGTCGGATGCCTGAACCTATGGATGCTTAACCAGCAGCTGAGTGAGTGTGAAAGTTCCCTGCTGACCCCTGGGTTGCTCAGGTGGCCTGTGCAGGACCCTGGCTCAGCTCTCAAGCTCTGCAGAGTGGGGTGGTGGAGATGCAGGGGAGGGGAAGGGAACTGCCATCTGAGCGCCAGCCTCCTGCTAGGCAAACCCGCCAGGGATGCTTGGAAGTGCTTTAATCTACACTGCTACAGCAGTGTGAGGTCTGGGGATTTGAATGGGGGCGGGGGAGGGATGTAAAAACCATAGCGCAGGATTTGGAAGGTCTGGTACAGGAGGAGAAAGCCCAGTTCCCTGTGCAGTCTGTTAGCCTGCTGCTCCACAATACTCTCTAGTTTCTATCCTTATTAGGTGATGGGAAGCACGCACTGCTAGAGTGCCCATTTGGGACAGGTATGACAGAAGTACCCTAATGTATCCAAGGACCCGCTTCTTCCTGTGACAGTCATCATCGTGGATGTATCTACTGAAGTCCTTTTAGAACCTGGGAGTGCTACTCAGCCTGCGTGGGAGTCCAGCTACGAGGTTCAGGTCCCCATTGGAGTGGGCAGCAAAAGGTTGTAGGCTGGAGTTCAGGTATTAAAGAGGTCGTGATGTCAAACTCAGGTTTGCTCACATTCTGGACGAATTCACCCTCTCTGTATCCTTACCCCACCCCCACTCCCACTCCTACCCGGTTCCTCAGCAATGACCTCAAAGACAGGGAGTGGACTGCTGCCTCCCTCCTGCAGAAGTGTAAGTAGCTCCAGCTATGACGTTATGGAAGCTCGGTCAGAGCTCTGATTGGTGGAAGAGCTACTGCGGACCCCCCACACCCTCAAGATCGAGAATAAGAGACCACGGCTCTGGGGACAAGACGCCTACAGCCGTGTGTGTGTGCTCTTCTGCAGTGGACACTTCACTCTCGCTGTTCCAACACGGGCTAGCAGGTGAGAAACTTAACTTCTCAACGCCTACAGCTCTCTCTCCTTTAGTTTGTTTCCTTTTGGTTGCTCTTTTTAATGCAGTATTTCACACTGTAGTCTAAGTTGGCCTGGCACCCACTATATAGCTCAAGTTGGTTCGTCTCAAACGCTAAGGTTCTCCTTTCTCAGCATCCTAAGGATGCGGATTACAGGCGCAAGCCACCACACCCCACTCTACTTGGATCCCTTTGCTGTCCTGGTTCCTTATCATTCCACATACATTTCCGCCTTCCTGCAGCCATTGTCAGAAAGTACAGTCTTGACATTTTCTTTTAACTAAAGTAAGTGGGACCCCTACGACTACTCAGCAGCACTGGAAGCTGGGCGACCCTATCTAGGCGCGTCTGTGCCCCCTCCTTGAGGGAAGGTGGTCTTGCCGCATCCTAACCAGTTTTAGGTTAAAGAGTTCCTTGGATTCGGGATTGGGGAGCACTGATCTTTTCTCTCAGATGTTTCCAGCCTTTAGCCTCCTGGGGTTATCAGCAAGCAGGTGGGTCTCGCTTCGCTGTGGGGAGGAGGAGTCCTCATCTGCGGTTCTGAGGTAGTTTAAAAAAAAAAATCTCCCAACTCTGCAGATGGAGAGAGGGGGATTAGTTCCAAGTTAACTTTCTTCCCCAGGGCAACCTCTCAGAAAGGGTGATTATAATAATTTCAACCTGTTAGAAATCCTTAGCAGCGGGACAGCAAGGCGCAGGGATCTCTGGGTGGTTTTTGGTTTCTTCACAGATGAGAGCCAAAGGGGCGCGGCACGTGTGTTCTCCTGCAAGCTGGGGGCAAATGAGTGCCGGTAGCTCCTCCTTGTTCTTAAACCGAGCAGAAACTGCAAACCACATCTACTCTCCCCCACTCGTTTCTGCTCTATCAAGCCACTCACCACACTGCATCTACTGCACGTTTTGAGAGCTGCAGTGTGGTAGGAGAAATAGAACCTGGGTCTATAGTCCTGAGCAATTGGACCATTCTTCTCTTCTTACAGAGACATCTTAATTAACTAGTGCGGCCGCCACCTCTAGAGGATCC 3’

The vector pAM-CGRP-FLPo-HA was used for initial testing of the CGRP promoter. DNA was de-novo synthesised containing the entire 2Kb CGRP promoter region upstream from FLPo, a 2A self-cleaving peptide sequence and 3x HA tags. This entire sequence was cloned into a vector containing AAV2 inverted terminal repeats (ITRs) (Murray et al., 2011) via KpnI and HindIII restriction sites.

The CGRP and cre-conditional FLPo or GFP (pAM-CGRP-Flex-FLPo or pAM-CGRP-Flex-GFP) were generated by replacing the CAGGS promoter in pAM-Flex-Empty (a cre-conditional vector containing AAV2 ITRs; Murray et al., 2011) with the CGRP short promoter via KpnI and XbaI restriction sites. FLPo or GFP was cloned in reverse orientation into the multiple cloning site using EcoRI and XhoI restriction sites.

### Generation of recombinant AAVs

AAV vectors used in this study were all packaged according to McClure et al., 2011 with some minor modifications. Briefly, human embryonic kidney (HEK) 293 cells were transfected via either calcium phosphate or Turbofect (Thermo Fisher Scientific) with individual AAV backbone plasmids as well as a 1:1 ratio of AAV1 (pH21) and AAV2 (pRV1) or AAVDJ helper capsid proteins and adenovirus helper plasmid pFdelta6. 48 hours post-transfection, the cells were harvested and AAVs purified using 1mL HiTrap heparin columns (Sigma), concentrated using Amicon Ultra centrifugal filter devices (Millipore) and purified using an AAVpro Purification Kit (Takara Bio).

The recombinant AAVs we designed and produced for selective EVN targeting were, in order of experimental use:

1. AAV[2/1]-CGRP-FLPo-HA
2. AAV[DJ]-CGRP-FLEx-GFP
3. AAV[DJ]-CGRP-FLEx-FLPo

Other AAV vectors used in this study were:

- AAV[DJ]-Ef1a-frt-H2BG-TVA (for rabies monosynaptic tracing; Reardon et al., 2016)
- AAV[DJ]-Syn-frt-H2BG-N2cG (for rabies monosynaptic tracing; Reardon et al., 2016)
- AAV[DJ]-EF1a-fDIO-TeLC-GFP (for behavioural recordings after blocking EVN transmission; ETH Zurich Vector Core)
- AAV[DJ]-EF1a-fDIO-GCaMP6s (control for behavioural recordings with TeLC injections; Addgene plasmid #105714)

### Animals

All animal experiments were performed under UK Home Office license (PPL: PE4FA53CB) according to the United Kingdom Animals (Scientific Procedures) Act 1986. Male and female mice aged between 8-16 weeks old from C57BL/6J and B6J.ChAT-IRES-Cre::Δneo (ChAT-Cre; JAX stock no.: 031661; Rossi et al., 2011) homozygous animals were used for experiments.

Initial experiments used ChAT-cre::tdTomato mice that were generated in house from a cross between homozygous strains B6J.ChAT-IRES-Cre::Δneo and ROSA-loxP-STOP-loxP-tdTomato (Madisen et al., 2010). These mice expressed tdTomato exclusively in cholinergic neurons and were used to demonstrate the location of EVN neurons (ChAT-positive) with peripheral fluoro-gold labelling.

All animals were housed in temperature-controlled environment with a 12hr light/dark cycle (red lights on at 0900) and free access to food and water. Surgeries and behavioural experiments were carried out during the dark phase of the cycle. All efforts were made to minimise animal suffering.

### Surgical procedures

#### Stereotaxic injections

Stereotaxic injections into the EVN were made as described previously (Murray et al., 2011), adapted from Cetin et al. (2006). Briefly, mice were anesthetised with isoflurane in oxygen (4% induction; 2% maintenance during surgery) and given a subcutaneous injection of analgesics (Metacam at 5mg/kg). The mouse was fixed in a stereotaxic frame (Model 942, Kopf, USA) and an incision was made in the skin such that bregma and lambda could be exposed and visualised. A small burr hole was made above the injection site (relative to bregma) where viruses were injected using a pulled 3.5” borosilicate glass capillary with 8µm bore width (Cat. no.: 3-000-203-G/X; Drummond, USA) and Nanoject II or Nanoject III (Cat. No.: 3-000-207 and 3-000-204 respectively; Drummond, USA). Stereotaxic coordinates relative to bregma for specifically targeting the EVN were as follows: anterior/posterior -5.80mm; lateral +0.60mm; depth from brain surface -4.4 and -4.3mm. Each depth received 100nl of AAVs or 200nl of rabies (CVSN2c-ΔG-EnvA-mCherry, Sainsbury Wellcome Centre Virology Core Facility) depending on the surgery. Following viral injection, the skin was closed with sutures (Cat. No.: VR493; Ethicon, USA).

Subsequent rabies injection or histology began no sooner than 14 days after the initial injection of AAVs to allow sufficient time for their expression. Likewise, histology following rabies injections also took place 14 days after rabies injection for the same reason.

#### Retrograde labelling

To identify EVN neurons fluoro-gold was injected into the right horizontal semi-circular canal of ChAT-Cre::tdTomato or Chat-Cre mice. These retrograde injections were made as described previously (Mathews et al., 2015). Briefly, mice were anaesthetised with 4% isoflurane in oxygen, given appropriate analgesics and maintained on 2% isoflurane in oxygen during surgery. The area behind the ear was shaved and cleaned, and an incision was made 1mm behind the right pinna. The muscles were separated such that the horizontal semi-circular canal was visualised. A 23G needle was used to thin the bone of the horizontal semi-circular canal until a small hole (approximately 100µm) was made. A 29G needle attached to a 1ml insulin syringe was used to carefully introduce 100nl of 2% fluoro-gold (Sigma-Aldrich) in saline into horizontal canal and the skin sutured. Animals were sacrificed 3 days later by transcardial perfusion described below.

### Tissue preparation and histology

Mice were deeply anaesthetised with intra-peritoneal injection of pentobarbital (200mg/ml concentration at 10mg/kg dosage; Dolethal) and subsequently transcardially perfused with 0.1M phosphate buffer saline (PBS, pH 7.4) followed by 4% paraformaldehyde in 0.1M PBS. The brains were harvested and immersion-fixed in 4% paraformaldehyde in 0.1M PBS for a minimum of 2hr at room temperature. 50µm thick coronal brain sections were sliced on a Leica V1200 vibratome and collected and stored in tissue culture wells filled with 0.1M PBS and 0.01% sodium azide solution until use.

Free-floating sections were incubated in primary antibodies diluted in 0.1M PBT (0.1M PBS with 0.1% Triton X-100 and 1% BSA) overnight. They were then washed three time in 0.1M PBS for 10 mins each before secondary antibody incubation (diluted in 0.1M PBS) for 2hrs. After washing again three times with 0.1M PBS for 10mins each, the sections were mounted onto Superfrost+ (Thermo Scientific) glass slides, dried and cover-slipped with DAPI-Fluoromount (SouthernBiotech 0100-20). Primary and secondary antibodies used for different AAV and rabies surgeries are summarised in Table 1. All sections were imaged using the Zeiss AxioScan Z1 slide scanner or Zeiss AxioImager at 10x or 20x magnification.

**Table 1.** Antibodies used throughout for immunohistochemistry experiments. Dilutions and catalogue number provided.

| Experiments | Primary Antibody<br>(dilution, catalogue number) | Secondary Antibody<br>(dilution, catalogue number) |
| --- | --- | --- |
| *Figure 1* | Chicken anti-GFP (1:1000, AB13970) | Anti-Chicken 488 (1:1000, A11039) |
|  | Rabbit anti-FG (1:200, AB153-I) | Anti-Rabbit 594 (1:1000, A21207) |
| *Figure 2* | Mouse anti-mCherry (1:1000, AB125096) | Anti-Mouse 594 (1:1000, A21203) |
|  | Sheep anti-ChAT (1:250, AB18736) | Anti-Sheep 647 (1:1000, A21448) |
|  | Chicken anti-GFP (1:1000, AB13970) | Anti-Chicken 488 (1:1000, A11039) |
| *Figure 3* | Rabbit anti-TeLC (1:1000, AB53829) | Anti-Rabbit 647 (1:1000, A21245) |
|  | Chicken anti-GFP (1:1000, AB13970) | Anti-Chicken 488 (1:1000, A11039) |

### Cell counting

#### Manual

For monosynaptic rabies tracing, the cell numbers of ‘starter’ EVN neurons (defined as expressing both the nuclear GFP from the AAV and mCherry from the RABV) was determined by eye, focusing through 50 µm thick coronal sections containing the EVN and counting the nucleoli identified with DAPI staining in consecutive sections containing the EVN.

#### QUINT workflow

In rabies tracing experiments where manual cell counting was not practical, cells were quantified and spatially analysed in a semi-automated manner using open-source software in QUINT workflow (detailed in Yates et al., 2019). Briefly, serial sections were registered to a 3D reference atlas (Allen Brain Mouse Atlas 2017) using QuickNII software. These sections are then segmented into cell, tissue and background using ilastik software. Finally, Nutil software was used to merge custom atlas maps from QuickNII with segmented images from ilastik to quantify cells and their coordinates. These data were exported to Microsoft Excel for further analysis. All output cells and their locations were cross-checked against original JPEG images of sections to ensure accurate counting and location reporting – here, mislabelling was manually adjusted. Importantly, the Allen Brain Mouse Atlas 2017 used in QuickNII does not include the location for the EVN and so EVN cells were misplaced as medial vestibular nucleus neurons. These were manually adjusted in post-processing.

### Behaviour recording and analysis

Behavioural assessment of TeLC expressing and control mice behavioural recordings took place at 7, 10 and 14 days post-AAV surgeries (see vectors listed above for details).

#### Open field

Mice were allowed to freely explore a square white Perspex arena for a period of 5 minutes while being recorded by an overhead Doric Behaviour Tracking video camera recording at 30 frames per second. To avoid effects of habituation on experiments mice were exposed to the arena for 5 minutes on the three days prior to experimental recordings.

Videos were analysed in EthoVision XT (Noldus) video tracking software. Here, centre-point tracking feature was used to track and generate total distance travelled as well as heat maps of movement where pixel colour represents total time spent at the location (more time spent corresponds to lighter blue colour). Distance travel data was exported to Microsoft Excel for plotting and further analysis. TeLC and control conditions were compared using student’s paired T-test.

#### Balance beam

The balance beam test (adapted from Tung et al., 2014) was used to measure subtle motor changes in mice where EVN transmission was blocked. The beam (1cm square; 83cm in length) was elevated and suspended between two platforms of different heights (13cm on the left and 19cm on the right) to create a slight incline of 3 degrees. On the right-hand side, the platform included a stage for mice to walk to and rest. A mirror was placed below and angled so to reflect the underside of the beam and mice onto the video recorder. Mice were habituated and trained to walk on the beam prior to AAV injections by placing the mouse progressively further away from the end platform and stage. Video recording from the right-side view of the mouse was carried out with a high-speed camera (Ximea) recording at 200 frames per second.

Video were analysed in MaxTraq Software (Innovision Systems) for speed of traversing across the beam and the nose to tail angle of each mouse measured at the centre of the beam. Tracking data was exported to Microsoft Excel for plotting and further analysis. TeLC and control conditions were compared using student’s paired t-test.

## Results

### Molecular targeting of EVN neurons

EVN neurons are known to co-localise ChAT and CGRP in mice (Ohno et al., 1991; Mathews et al., 2015). We used this information to design an intersectional genetic strategy to selectively target these neurons via a stereotaxic injection into the brainstem. First, we anatomically visualised EVN neurons, which are known to project and synapse on to vestibular hair cell receptors and primary afferents in the peripheral sensory apparatus, by injecting the retrograde tracer fluoro-gold into the horizontal semi-circular canal of ChAT-cre::tdTomato mice.

Fluoro-gold labelling of EVN neurons, which are located dorsolateral to the genu of the seventh cranial nerve, was observed overlapping with ChAT-positive putative EVN neurons, confirming that EVN neurons can be targeted using cholinergic labelling in ChAT-Cre animals (Figure 1A). However, cholinergic neurons are also observed in the nearby abducens nucleus limiting the ability to target the EVN with stereotaxic injections alone. We therefore attempted to further refine this targeting by complimenting the cre line with an AAV that only expressed a transgene in neurons that were both ChAT (cre) and CGRP positive. This AAV used a CGRP promotor fragment to express a nuclear tagged human influenza hemagglutinin (HA) in CGRP-positive cells. We used this in combination with a Cre-dependent AAV to express GFP in ChAT-positive cells in ChAT-Cre mice. Stereotaxic co-injections of both AAVs in ChAT-Cre mice allowed us to visualise ChAT-positive cells in green and CGRP-positive cells in red. We observed co-localisation of GFP and HA in EVN neurons (Figure 1B).

**Figure 1.**
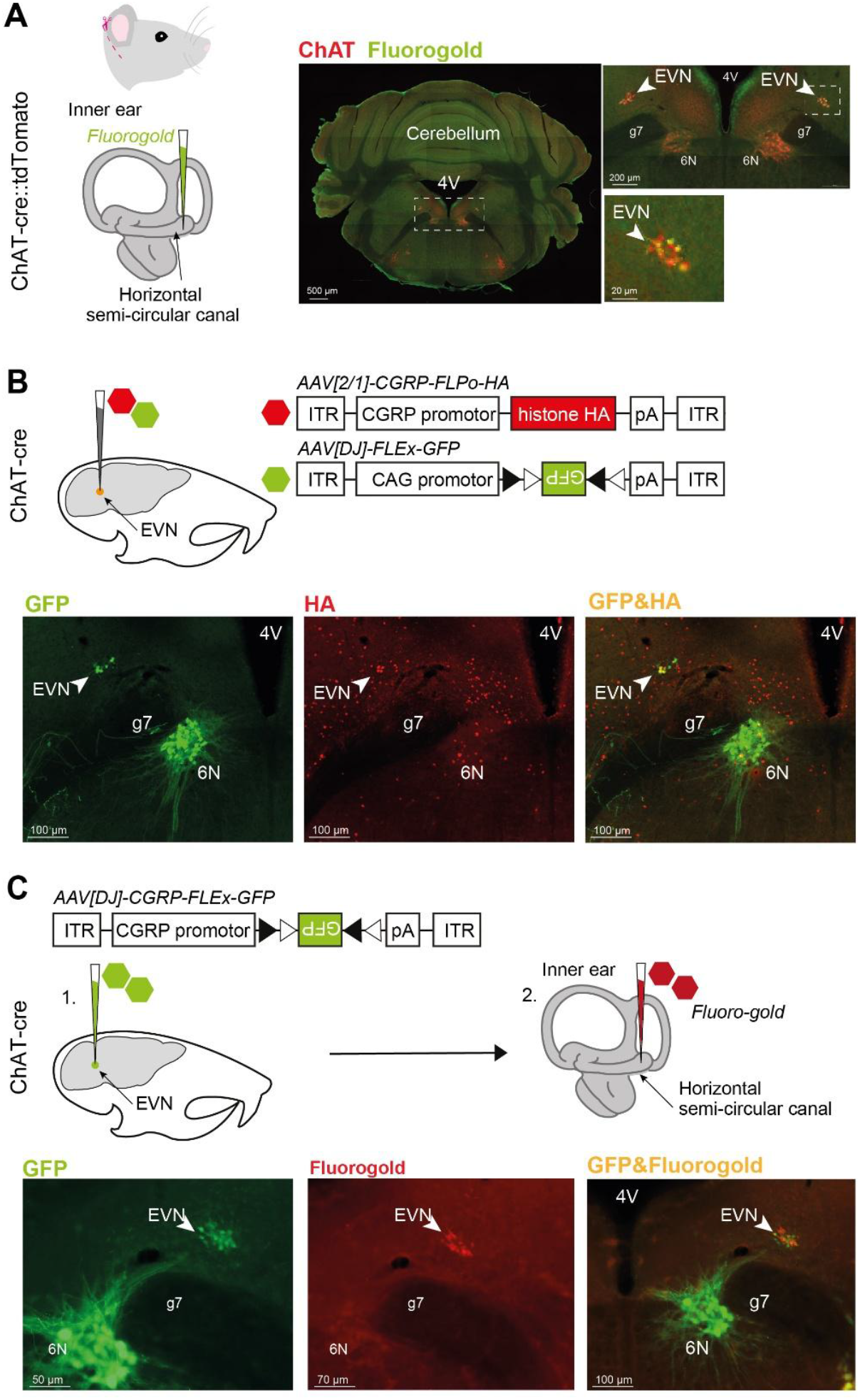
Molecular-genetic targeting of EVN neurons. *(A)* The fluorescent retrograde tracer fluoro-gold (FG) was injected into the horizontal semi-circular canal of ChAT-cre::tdTomato mice. Gold FG staining overlaps with red ChAT-positive, putative EVN neurons in red (arrow, insets), confirming EVN is located dorsolateral to the genu of the seventh cranial nerve (g7), and can be targeted with ChAT-cre animals. *(B)* In ChAT-cre mice, a 1:1 ratio of cre-dependent AAV expressing GFP and a HA-expressing AAV (red) under control of the CGRP promotor was stereotaxically injected into the EVN. Yellow overlap (arrow, far right image) confirms that CGRP and ChAT, can be used to selectively target EVN neurons. *(C)*. Combined strategy where a single AAV combines the CGRP promoter and a cre-dependent FLEX switch can be used to target EVN neurons. Confirmed by stereotaxic injection of this AAV into the EVN and fluoro-gold injection into the semicircular canal. *Abbreviations*: *4V* fourth ventricle; *CAG* chicken beta-actin; *Cb* cerebellum; *CGRP* calcitonin gene related peptide; *ChAT* choline acetyltransferase; *ITR* inverted terminal repeats; *pA* poly A; *VN* vestibular nuclei. Mouse inner ear schematic adapted from Schutz *et al*. (2014).

Next, we developed a cre-dependent AAV using this CGRP promotor to selectively express GFP in cells that are both ChAT and CGRP positive. Stereotaxic injections of this AAV allowed us to visualise GFP in EVN neurons. To ensure the AAV effectively targeted EVN neurons, we performed fluoro-gold injections in the horizontal semi-circular canal. Co-localisation of GFP with fluoro-gold was observed on average (± standard deviation) in 15±4 neurons (n=5 animals), representing approximately 40% of the nucleus, confirming this strategy selectively targets EVN neuron (Figure 1C).

### Monosynaptic inputs to EVN neurons

Selective expression of recombinases, such as flp recombinase, in genetically defined neurons provides a means to interrogate both their anatomy and function (Nectow and Nestler, 2020). Our intersectional approach provided a means to selectively target EVN neurons, and therefore we developed a viral vector (pAAV-CGRP-FLEx-FLPo) that combined the short CGRP-promoter and cre-conditional expression to drive flp-recombinase only in EVN neurons (Figure 2A). We first used this tool to identify direct synaptic inputs to EVN neurons.

**Figure 2.**
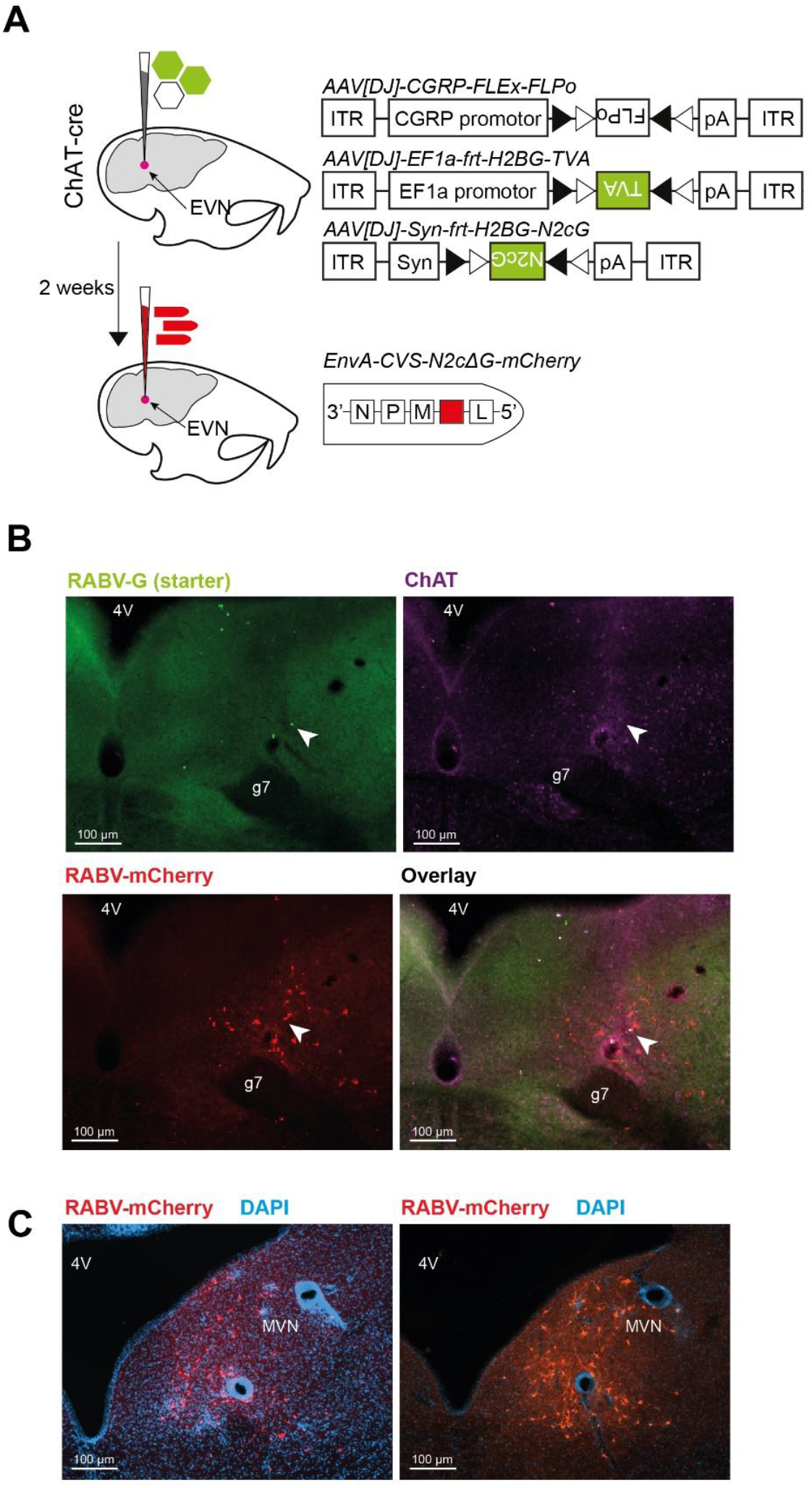
Monosynaptic rabies tracing of EVN neuronal inputs. *(A)* Strategy for rabies tracing in ChAT-cre mice where first avian receptor TVA (AAV[DJ]-EF1a-frt-H2BG-TVA) and rabies N2c glycoprotein (AAV[DJ]-Syn-frt-H2BG-N2cG) were stereotaxically injected with AAV[DJ]-CGRP-FLEx-FLPo in a 1:1:1 ratio to prime neurons for rabies tracing. Two weeks following initial rAAV injections, EnvA-psudotyped, glycoprotein-deficient rabies tagged with red fluorescent protein (EnvA-CVS-N2cΔG-mCherry) was stereotaxically injected to the same coordinates. *(B)* Successful transduction (arrow, overlay) of rabies G (arrow, green), and rabies virus (arrow, red) in a ChAT-positive EVN neuron (arrow, violet). *(C)* Surrounding monosynaptic EVN inputs from the MVN shown in red.

We co-injected a mixture of three AAVs into ChAT::Cre mice – AAV[DJ]-CGRP-FLEx-FLPo to drive selective FLPo expression only in EVN neurons, and AAV[DJ]-Ef1a-frt-H2BG-TVA and AAV[DJ]-Syn-frt-H2BG-N2cG to express the TVA receptor and rabies glycoprotein respectively in a Flp-recombinase dependent manner. The TVA receptor permits selective cell entry of EnvA-pseudotyped rabies virus, while rabies glycoprotein expression enables monosynaptic transfer of rabies virus (Reardon et al., 2016). 2 weeks following this AAV injection into the region of the EVN, we injected RABV-N2c-EnvA-mCherry at the same stereotaxic coordinates to target EVN neurons and identify their monosynaptic inputs (Figure 2B).

Almost all monosynaptic inputs to EVN neurons were found within the brainstem/midbrain region (such as vestibular nuclei, reticular formation, medullary and pontine regions and autonomic /arousal nuclei), suggesting modulation by local circuits (Figure 2C). No neurons were observed in cortical or thalamic regions, or in the spinal cord (data not shown).

Confirming the selectivity of our strategy we identified only five starter EVN neurons across three mice where the mixture of three AAVs co-localised with rabies following injections, demonstrating that the identified inputs came only from the EVN. These starter neurons were abundantly innervated from input regions (starter to input neuron ratio of 1:356). A categorised summary of direct monosynaptic inputs, all ipsilateral, to these neurons is provided in Table 2. Approximately 50% of the inputs to EVN neurons came from the neighbouring medial vestibular nucleus (MVN). Self-innervation by other EVN neurons accounted for ∼7% of inputs, while other vestibular nuclei, including the lateral vestibular nucleus (LVN) and spinal vestibular nucleus also contributed inputs (1% and 3% respectively). Other movement associated areas such as the pontine reticular nucleus and sublaterodorsal nucleus each provided ∼3% of EVN inputs. Eye movement associated areas nucleus prepositus and abducens nucleus also provided direct EVN innervation.

**Table 2.**
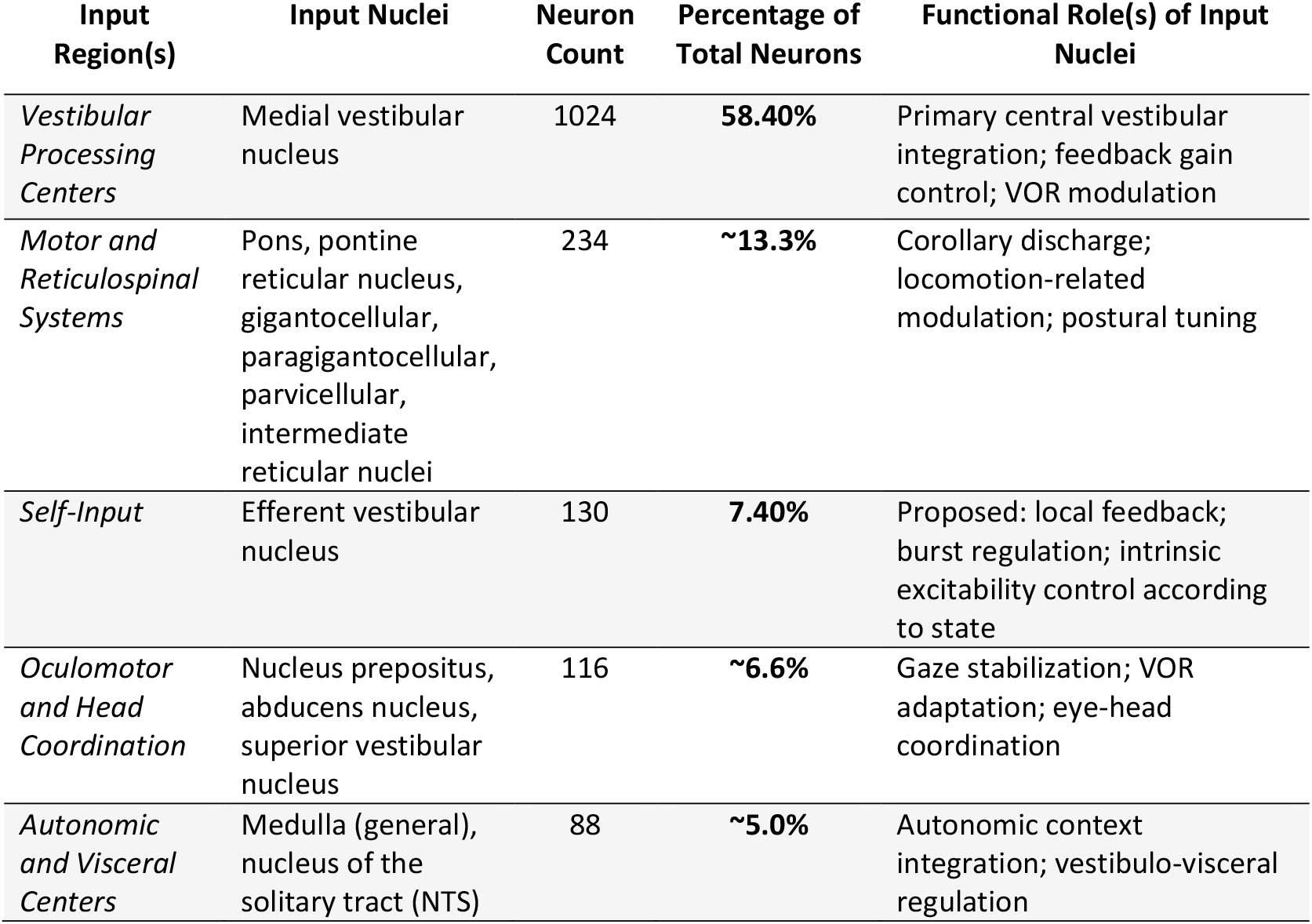
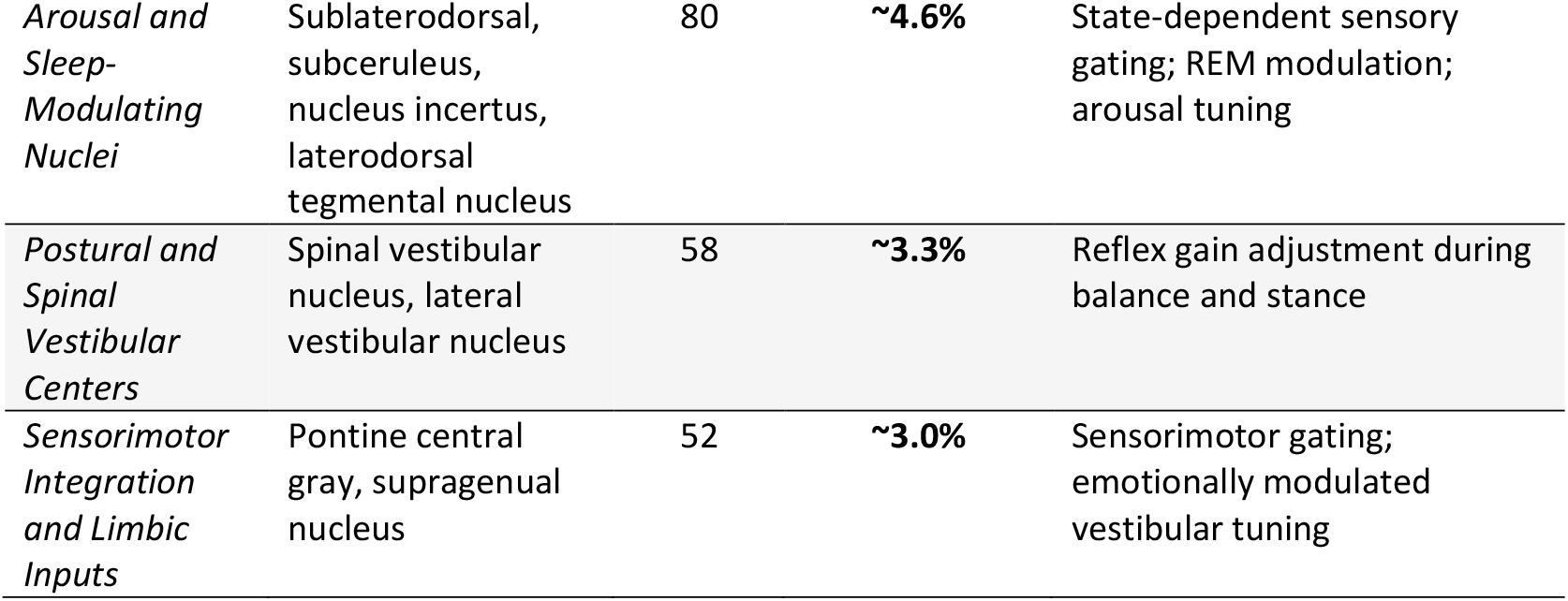
Monosynaptic inputs to the efferent vestibular nucleus (EVN) organized by brain regions. The novel intersectional rabies tracing described in this paper was used to determine the direct inputs to the EVN. Input nuclei neuron counting and atlas mapping and overlay was done using QUINT workflow and open-source software (Yates et al., 2019). Results are categorized into brain regions in order of percentage of total inputs. Broad functional roles of input nuclei also provided. Inputs were ipsilateral from five starter EVN cells across three mice.

| Input Region(s) | Input Nuclei | Neuron Count | Percentage of Total Neurons | Functional Role(s) of Input Nuclei |
| --- | --- | --- | --- | --- |
| *Vestibular Processing Centers* | Medial vestibular nucleus | 1024 | **58.40%** | Primary central vestibular integration; feedback gain control; VOR modulation |
| *Motor and Reticulospinal Systems* | Pons, pontine reticular nucleus, gigantocellular, paragigantocellular, parvicellular, intermediate reticular nuclei | 234 | **~13.3%** | Corollary discharge; locomotion-related modulation; postural tuning |
| *Self-Input* | Efferent vestibular nucleus | 130 | **7.40%** | Proposed: local feedback; burst regulation; intrinsic excitability control according to state |
| *Oculomotor and Head Coordination* | Nucleus prepositus, abducens nucleus, superior vestibular nucleus | 116 | **~6.6%** | Gaze stabilization; VOR adaptation; eye-head coordination |
| *Autonomic and Visceral Centers* | Medulla (general), nucleus of the solitary tract (NTS) | 88 | **~5.0%** | Autonomic context integration; vestibulo-visceral regulation |
| *Arousal and Sleep-Modulating Nuclei* | Sublaterodorsal, subceruleus, nucleus incertus, laterodorsal tegmental nucleus | 80 | ~4.6% | State-dependent sensory gating; REM modulation; arousal tuning |
| *Postural and Spinal Vestibular Centers* | Spinal vestibular nucleus, lateral vestibular nucleus | 58 | ~3.3% | Reflex gain adjustment during balance and stance |
| *Sensorimotor Integration and Limbic Inputs* | Pontine central gray, supragenual nucleus | 52 | ~3.0% | Sensorimotor gating; emotionally modulated vestibular tuning |

### EVN inhibition in behaviour

A complete understanding of the function of any neural circuit requires the ability to selectively alter the synaptic output of that group of neurons. To demonstrate whether our novel method could also be used to block synaptic output from EVN neurons we combined the selective flp-recombinase expression described above with an AAV that drives the expression of tetanus toxin light chain (TeLC) in a flp-dependent manner (Figure 3A).

**Figure 3.**
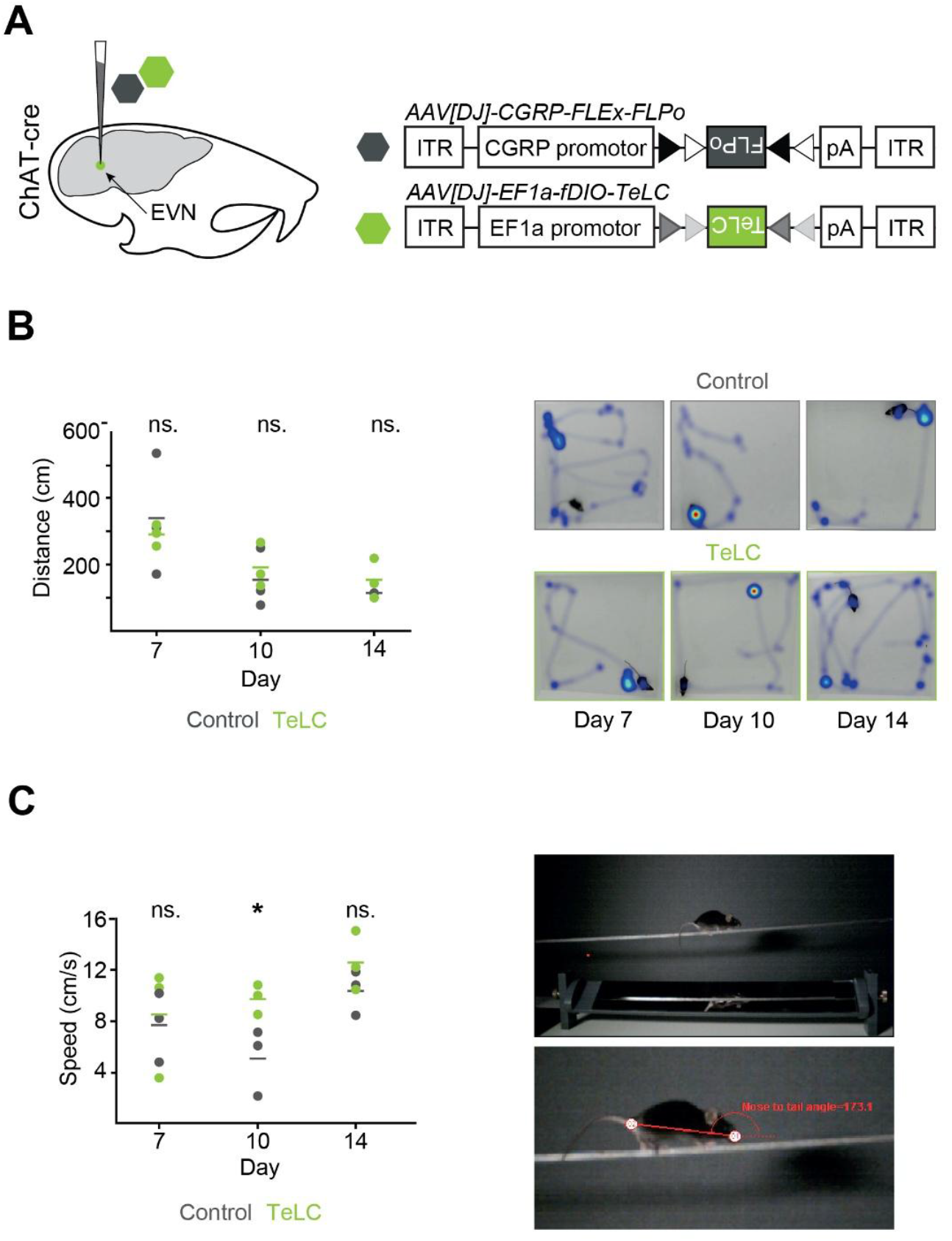
Selective disruption of EVN neurotransmission and behavioural assays. *(A)* Tetanus toxic light chain (TeLC) was selectively expressed in EVN neurons to block their transmission in ChAT-Cre mice via stereotaxic co-injection of AAV[DJ]-CGRP-FLEx-FLPo and AAV[DJ]-EF1a-fDIO-TeLC-GFP in a 1:1 ratio (n=3). In control mice, TeLC was replaced with TeLC with AAV[DJ]-EF1a-fDIO-GCaMP6s (n=3). *(B)* No significant difference was observed across control and TeLC expressing mice in an open field (white Perspex box) arena in the first 60 seconds of recordings, measured on days 7, 10 and 14 following stereotaxic surgery. *(C)* TeLC expressing mice demonstrated a significantly faster time to traverse the balance beam than controls on day 10 following stereotaxic surgery in line with peak TeLC activity and exposure. ^*^p<0.05.

TeLC is a potent means to block synaptic transmission and does so via the cleavage of the synaptic protein VAMP2, preventing vesicle docking. AAV mediated expression of TeLC is known to completely block release of neurotransmitters with effects beginning ∼10 days after AAV injection (Murray et al., 2011).

In experimental animals we performed bilateral co-injections of our EVN targeting AAV (AAV[DJ]-CGRP-FLEx-FLPo) with the Flp-dependent AAV[DJ]-EF1a-fDIO-TeLC-GFP into ChAT-Cre mice to selectively block synaptic transmission only from EVN neurons. Control animals received an injection of AAV[DJ]-EF1a-fDIO-GCaMP6s instead of TeLC at the same coordinates.

To assess whether our technique could be used to interrogate the behavioural role of EVN neurons, we performed two preliminary behavioural assays. Prior to formal behavioural assessment we examined animals for any gross behavioural abnormalities, especially those associated with peripheral vestibular abnormalities (Hardisty-Hughes et al., 2010). This included tilting of the head and circling. We also examined mice for other phenotypes such as reduced grooming and weight loss, but we did not observe any changes in these metrics (data not shown).

For a preliminary assessment of motor abnormalities in animals with disrupted EVN function we used an open field test. Mice were tracked in an open field arena (n=3 per condition) for 5 minutes and their path length, movement velocity and turning velocity analysed at 7-, 10- and 14-days post injection. No difference was found between control animals and those lacking EVN output (Figure 3B), suggesting the EVN is not required for gross motor function.

For a behavioural assessment that would require more engagement of the vestibular system we used the balance beam test, where animals with a vestibular impairment show balance deficits (Tung et al., 2014). As above animals were tested 7-, 10- and 14-days post AAV injection. Both control and EVN-disrupted animals were able to traverse the balance beam following a training period. No difference was found between groups in terms of the number of foot slips made by each animal as they walked across the beam, with both groups of animals making minimal slips per trial (foot slips per trial – day 7: control 0.4±0.4, TeLC 0.5±0.2; day 10: control 1±0.4, TeLC 0± 0; day 14: control 0.5±0.2, TeLC 0.17±0.17).

Animals with a disrupted EVN were consistently faster at traversing the balance beam (speed (cm/s) – day 7: control 6.8±1.1, TeLC 8.8±1.3; day 10: control 6.0±1.3, TeLC 10.4±0.9; day 14: control 8.6±1.0, TeLC 11.8±1.7). This reached statistical significance at 10 days following injection (Student’s t-test p=0.021). No other differences were found between control and EVN disrupted groups.

## Discussion

It is well accepted that efferent vestibular nucleus (EVN) neurons contribute to fine-tuning of vestibular sensory input by modulating firing patterns, signal timing and sensitivity of vestibular afferents and hair cells (reviewed in Mathews et al., 2020). However, the behavioural role of the EVS continues to be debated. An inability to target the EVN selectively has meant that probing its function in awake behaving animals has not been achievable. Here, we add to the toolkit for studying efferent vestibular system (EVS) function with a novel AAV-based EVN-targeting virus that draws on recent technological advances in neuroscience. Modern systems neuroscience relies on the linking of genetically defined neuronal subtypes with both their place in the wider neuronal architecture and animal behaviour. Much initial progress has been made by categorising neurons based on expression of single genes (Callaway, 2005). However, intersectional genetics, where the expression of two or more genes is used to define and target a neuronal population, is becoming more important as our understanding of neuronal diversity increases (Dymecki et al., 2010). We apply these principles to develop a novel molecular genetic tool for use in EVS research.

We combined a cre-recombinase driver line (ChAT::Cre) with a short AAV promoter active in CGRP-positive neurons, to drive selective gene expression only in neurons which express both these genes. As EVN neurons are one of the few neuronal subtypes to co-express these genes in the brainstem, stereotaxic injection permitted their selective targeting. We used this intersectional strategy to express a separate recombinase, FLPo, selectively in EVN neurons. Paired with recombinase-dependent AAVs, this allows for the selective anatomical mapping and activity manipulation of EVN neurons. We used this to assess 1) direct (monosynaptic) EVN inputs, and 2) perform preliminary behavioural assessment of EVS function. Though our anatomical and behavioural studies restricted genetic interventions to modified rabies tracing and expression of TeLC, this same system can be used for other systems neuroscience interventions such as the selective expression of fluorescent proteins, or optogenetic or chemogenetic modulators (Nectow and Nestler, 2020) for further direct interrogation of EVS function.

### 1) Direct synaptic inputs indicate EVS function as multimodal processing center

Mapping circuit architecture to function has become an important means to elucidate nervous system function. We know the EVN is sensitive to a heterogeneous input profile (Mathews et al., 2015) from a variety of brain regions (Zhou et al., 2018, Metts et al., 2006). So, we reasoned that selectively targeting and mapping the input circuitry for EVN neurons could provide behavioural clues and support a functional role for the EVS.

Selective targeting of EVN neurons allowed us to utilise the monosynaptic rabies system to assess direct inputs to EVN neurons. We show direct input from a highly specific subset of brainstem regions. The majority (58%) of all direct inputs originate from the medial vestibular nucleus (MVN) – which itself receives prominent vestibular afferent input – highlighting a strong functional coupling of central modulation on first-order vestibular processing in real time. We also observed input from other EVN neurons (7%) and smaller but functionally significant contributions from nuclei involved in eye movement control, posture, autonomic regulation, and behavioural state modulation.

Instead of a single explicit functional role, decades of research have either demonstrated directly or hypothesised EVN activation across a wide range of behavioural and physiological conditions. By quantitatively mapping the monosynaptic inputs to EVN neurons and integrating the known functions of these source regions, we infer that the EVN supports three distinct yet overlapping functional paradigms in the literature. The majority of EVN inputs support (i) vestibular plasticity and gaze stabilization (∼65%) while (ii) state-dependent gating (∼16%) and (iii) predictive motor suppression (∼13%) represent smaller but important functions. This inclusive framework accounts for the long-standing variability in EVN-linked behaviours and is consistent with the proposed EVN role as a dynamic, multimodal modulator of peripheral vestibular encoding.

#### I. Vestibular plasticity and gaze control

In addition to the MVN, the EVN receives input from oculomotor-associated nuclei, notably the abducens nucleus and nucleus prepositus, which are integral to horizontal gaze control and gaze holding, respectively. Collectively these regions integrate vestibular signals with motor commands for gaze stabilisation and in refining vestibulo-ocular reflex (VOR) gain and adaptation (Dietrich et al., 2017; McFarland and Fuchs, 1992; Mettens et al., 1994; Müri et al., 1996; Lisberger and Miles 1980). Their strong input indicates the EVN aligns peripheral vestibular encoding with centrally calibrated gaze shifts. In agreement with our anatomical data, early hypothesis of EVS function included anticipation of volitional head movement and the ensuing gaze shift (Goldberg and Fernàndez, 1980; Highstein, 1991; Brichta and Goldberg, 2000). Furthermore, the EVS has been suggested to have a role in vestibular plasticity, particularly regarding the vestibuloocular reflex (VOR) through signalling via alpha-9 nicotinic acetylcholine receptors (α9 nAChRs) expressed at efferent vestibular synapses on hair cells, which can elicit inhibitory responses in afferents (Elgoyhen et al., 1994; Hiel et al., 1996; Anderson et al., 1997; Holt et al., 2017; Zhou et al., 2013).

Separately, the EVS has also been implicated in motion sickness symptoms. CGRP expression levels increased in the EVN in rats with motion sickness (Xiaocheng et al., 2012), α9 nAChRs knockout mice with an attenuated EVN showed reduced motion sickness symptoms (Tu et al., 2017), and young adult mice displayed EVN activation (measured via cFos) during motion sickness symptoms (Lorincz et al., 2023). It is broadly accepted that motion sickness symptoms arise out of sensory mismatch and conflict between visual, proprioceptive and vestibular information. The EVN’s involvement is consistent with this model, particularly given its dense and selective innervation from brainstem and midbrain regions that integrate multimodal sensory information.

#### II. State-dependent sensory gating

Inputs from brainstem regions involved in arousal, visceral regulation, and motor control suggests that the EVN plays a key role in state-dependent sensory gating of vestibular input. Inputs from arousal- and sleep-related nuclei (e.g., sublaterodorsal, subceruleus, and nucleus incertus) imply that EVN activity may be modulated across sleep-wake cycles or during shifts in attentional state, potentially allowing the vestibular system to prioritize or suppress sensory input as needed (Boissard et al., 2003; Fraigne et al., 2015; Ryan et al., 2011; Valencia Garcia et al., 2017). Autonomic and visceral centers, including the nucleus of the solitary tract and medullary inputs, could provide interoceptive context enabling the EVN to modulate vestibular sensitivity during challenges like nausea, hypotension, or cardiovascular changes (Andresen and Paton, 2011; Zoccal et al., 2014). Meanwhile, inputs from sensorimotor integration regions (pontine gray and supragenual nuclei) and postural control centers (spinal and lateral vestibular nuclei) support the idea that the EVN dynamically calibrates vestibular afferent output based on ongoing motor activity, body and head orientation, and valence (Biazoli et al., 2006; Xiao et al., 2023). Previous studies have shown EVN activity increases during aroused states in toadfish (Highstein and Baker, 1985), and vestibular reflex sensitivity is enhanced under threatening postural conditions in humans (Lim et al., 2016; Naranjo et al., 2016). Multimodal stimulation such as light touch, sound, visual stimuli effectively increase EVN discharge (Highstein and Baker, 1985; Highstein, 1991; Münz and Claas, 1991; Roberts and Russell, 1972).

#### III. Predictive motor gating

Input from vestibulospinal and reticulospinal systems including the pontine reticular formation and lateral vestibular nucleus (which receives both motor input directly as well as through the cerebellum; Witts and Murray, 2019), supports a role in predictive motor gating of vestibular input. These areas are central to generating motor programs for locomotion, posture, and axial control (Takakusaki et al., 2016). Their input to the EVN could provide efference copy information about impending self-motion, enabling the EVN to adjust the gain of vestibular signals during volitional movement. In larval Xenopus and zebrafish, the EVS attenuates afferent discharge in phase with locomotor activity (Chagnaud et al., 2015; Lunsford et al., 2019), helping to suppress reafferent noise and maintain perceptual clarity during self-generated motion.

Our assessment of EVS inputs provides additional evidence for the potential behavioural roles of this nucleus. We propose that the EVN is an important component of reflexive, adaptable vestibular sensorimotor processing. Specifically, the EVN is a context-sensitive, integrative processor of vestibular, motor and internal state information for dynamically calibrating vestibular signalling in real time. This hypothesis is also consistent with the self-input we observed – the EVN likely uses an internal feedback system as an auto-tune mechanism when filtering diverse information.

### 2) Behavioural assessment of EVS function

We performed a preliminary behavioural assessment of EVS function via the selective expression of TeLC in EVN neurons. TeLC provides a potent block of synaptic transmission approximately 10 days after AAV injection (Murray et al., 2011). Importantly, animals were overtly normal and showed no health problems following EVN disruption. Additionally, animals showed no gross motor defects in the open field and were able to traverse a balance beam. This latter result potentially indicates that the EVS is not required for all peripheral vestibular function as this task is heavily dependent on the vestibular system and mice with vestibulopathies show clear deficits in this behaviour (Tung et al., 2014; Tung et al., 2016).

Mice with a disrupted EVN did show a mild phenotype of traversing the balance beam faster than control animals. This counterintuitive result (as you may expect that animals with a suboptimal vestibular system would be slower at a balance task) may indicate an inability of animals to regulate their own locomotor speed during challenging conditions, perhaps because of disrupted motor efference copy information (Chagnaud et al., 2015). This result also hints that the EVN normally acts to constrain movement speed, acting as a behavioural brake or stabilizer during dynamic motor tasks, possibly at the cost of long-term sensorimotor adaptability or precision. However, further behavioural assessment will be required to confirm this result.

One possible explanation for the limited behavioural effects observed could be the small number of cells ultimately affected by virally delivered manipulations. The EVN has few neurons (average 47 neurons in each hemisphere in mice; Mathews et al., 2015, Lorincz et al., 2021) and only transducing a proportion of the cells within it results in a small overall intervention. Our CGRP driven EVN targeting isolated on average fifteen EVN neurons, approximately one third of all EVN neurons. While this is useful for anatomical applications, such as rabies tracing, it is challenging when looking at behavioural assays where a complete inhibition of the whole EVN would provide more clarity in results. Additionally, given the highly plastic nature of the vestibular system and its ability to compensate for perturbations (Gittis and du Lac, 2006) the use of a chronic manipulation such as TeLC could obscure EVS function if the system were to compensate for this lack of output.

## Conclusions and next steps

Here, we provide a novel means of targeting and accessing the EVN that we use for monosynaptic tracing of inputs and behavioural assessments of EVS function. Our input mapping experiments utilise rabies tracing methodology and provide valuable new information of the anatomical organization of the efferent vestibular system. For the first time, we present a wholistic understanding of the type of information that the EVN receives and processes. Combined with the existing body of EVS research, our findings highlight the multimodal nature of the EVN, acting as a hub to integrate and update the vestibular periphery in real time with relevant internal and external information to support overall vestibular health.

Our molecular genetic strategy could also be used alongside bespoke behavioural tests to further demonstrate this filtering role or test alternative functional hypotheses. For example, head-fixed eye tracking for VOR performance or complex movement tasks such as narrower or curved beam experiments could be adapted to our technology. Our system is also compatible for use with acute circuit interventions such as chemo- or opto-genetics and calcium imaging which could provide a clearer understanding of EVS function.

EVS research continues, and the novel methodology presented here enables specific targeting of the EVN for functional research and, ultimately, faster progress in understanding this somewhat mysterious nucleus.

## Competing interests

The authors have no competing interests related to this work.

## Acknowledgements and funding

This work was supported by the Sainsbury Wellcome Centre core grant from the Gatsby Charitable Foundation and the Wellcome Foundation (090843/F/09/Z).

## Notes

### Competing Interest Statement

The authors have declared no competing interest.

